# Mechanism of membrane curvature generation and caveola formation by flat disc-like complexes of caveolin

**DOI:** 10.1101/2024.07.09.602763

**Authors:** Avishai Barnoy, Robert G. Parton, Michael M. Kozlov

## Abstract

Membrane sculpting by caveolin and accessory proteins is crucial for caveola formation. The discovery of the disc-like structure of caveolin oligomers has challenged earlier models for membrane shaping by caveolins. The flat shape of the caveolin discs apparently contradicts the large curvature of caveolar membranes these discs generate upon their predicted embedding into the cytoplasmic membrane leaflet. Here we have provided a mechanism for this phenomenon. We proposed that the central factor behind the membrane shaping by caveolin discs is a differential interaction of the membrane lipid monolayers with each other and with the hydrophobic faces of the caveolin discs. Based on this hypothesis we demonstrated by computations that the caveolin disc insertion causes elastic stresses of tilt and splay in the membrane monolayers, which, in turn, drive membrane kinking along the disc boundaries. The resulting membrane shapes are predicted to have a faceted appearance in agreement with observations and the estimated effective curvatures of these shapes are equal to those measured for caveolae in cells. We predicted and analysed the membrane-mediated repulsive forces developing between the inserted caveolin discs and discussed the strength of the counterforces needed to concentrate caveolin discs in the membrane plane. Besides recovering the major features of caveolar formation and morphology, our model provides a new mechanistic understanding of the role of cholesterol and other lipids with similar intrinsic curvature in the caveola assembly and control of caveolar size.

## Introduction

Caveolae are nanoscale invaginations of the plasma membrane which form a characteristic and abundant feature of the plasma membrane of many mammalian cell types ^1^. A typical caveola shape can be described as a 50-80nm diameter large nearly spherical bulb connected to the plasma membrane by a narrow neck ^1^.

Since their discovery more than 70 years ago ^2,3^, caveolae have remained a focus of cell biological research due to the constantly emerging new insights into their multiple functions in cell physiology ^4,5^, and the deepening understanding of the molecules and molecular complexes involved in caveola formation and dynamics ^6-8^. Yet, despite a wealth of phenomenological knowledge, the physical mechanisms governing caveola shaping by dedicated molecules remain elusive. The present work aims to propose such a mechanism by accounting for the most recent breakthrough in the structural biology of caveola-forming proteins along with the state-of-the-art information on the conformations of caveolar membranes and the role of lipids in caveola biogenesis.

Since detailed reviews are already available on the structure and function of caveolae ^6,7,9^, here we present only a sketch of the recently emerged experimental data relevant to the proposed model.

The two major protein families responsible for caveola formations are the integral membrane proteins, caveolins, and the peripheral filamentous proteins, cavins ^1,6^. These proteins self-assemble on the membrane surface into a dense two-layer complex, that has been proposed to have an intrinsically curved conformation and, therefore, sculpt the underlying membrane into the caveola bulb ^9^. Importantly, caveolins can generate caveola-like membrane invaginations by themselves in the absence of cavins. This was demonstrated by the expression of a wide range of recombinant caveolins in the heterologous host *Escherichia coli*, which resulted in the formation of membrane-enclosed vesicles, termed h-caveolae, derived from the cytoplasmic membrane^10^. The curvature, size, and caveolin density of h-caveolae were shown to be similar to those of native caveolae ^10^. This suggests that caveolins must be the primary factor in the mechanism of caveola formation with cavins playing auxiliary roles. Consistent with this, some invertebrate species have been shown to possess caveolin-dependent membrane invaginations in the absence of cavin proteins^6,11^.

The ability of caveolins to form the caveolar bulbs must be determined by the structure of the caveolin oligomer, serving as a minimal functional unit of membrane shaping, and the mode of this unit’s interaction with the membrane matrix. The data on these two issues was accumulated for the most ubiquitous member of the caveolin family, CAV1, referred to simply as caveolin, for brevity. Caveolin monomers are synthesized in the ER and then are transported to the Golgi complex where the minimal functional unit, the caveolin 8S complex, is assembled ^6,12^. The 8S complexes are believed to organize into higher order assemblies, the 70S complexes, in transit to the plasma membrane^12^. According to the recent experimental advance, a caveolin 8S complex consists of 11 tightly packed protomers arranged in such a way that the overall conformation of the complex is that of a flat disc of about 14nm diameter and 3nm thickness ^13,14^. One of the flat planes and the outer rim of a caveolin disc are hydrophobic, whereas the second plane is hydrophilic and contains both the amino- and carboxyl-termini of the protomers ^13^. This structure suggests a particular character of the caveolin discs’ association with the plasma membrane. It has been predicted that a caveolin disc embeds into the cytosolic membrane leaflet such that its flat hydrophobic plane comes into contact with the bottom surface of the extracellular membrane leaflet, the hydrophilic plane faces the cytoplasm, and about 250 lipid molecules of the cytosolic leaflet are pushed aside ^6,7,13-15^ (Fig.1).

**Figure 1.**
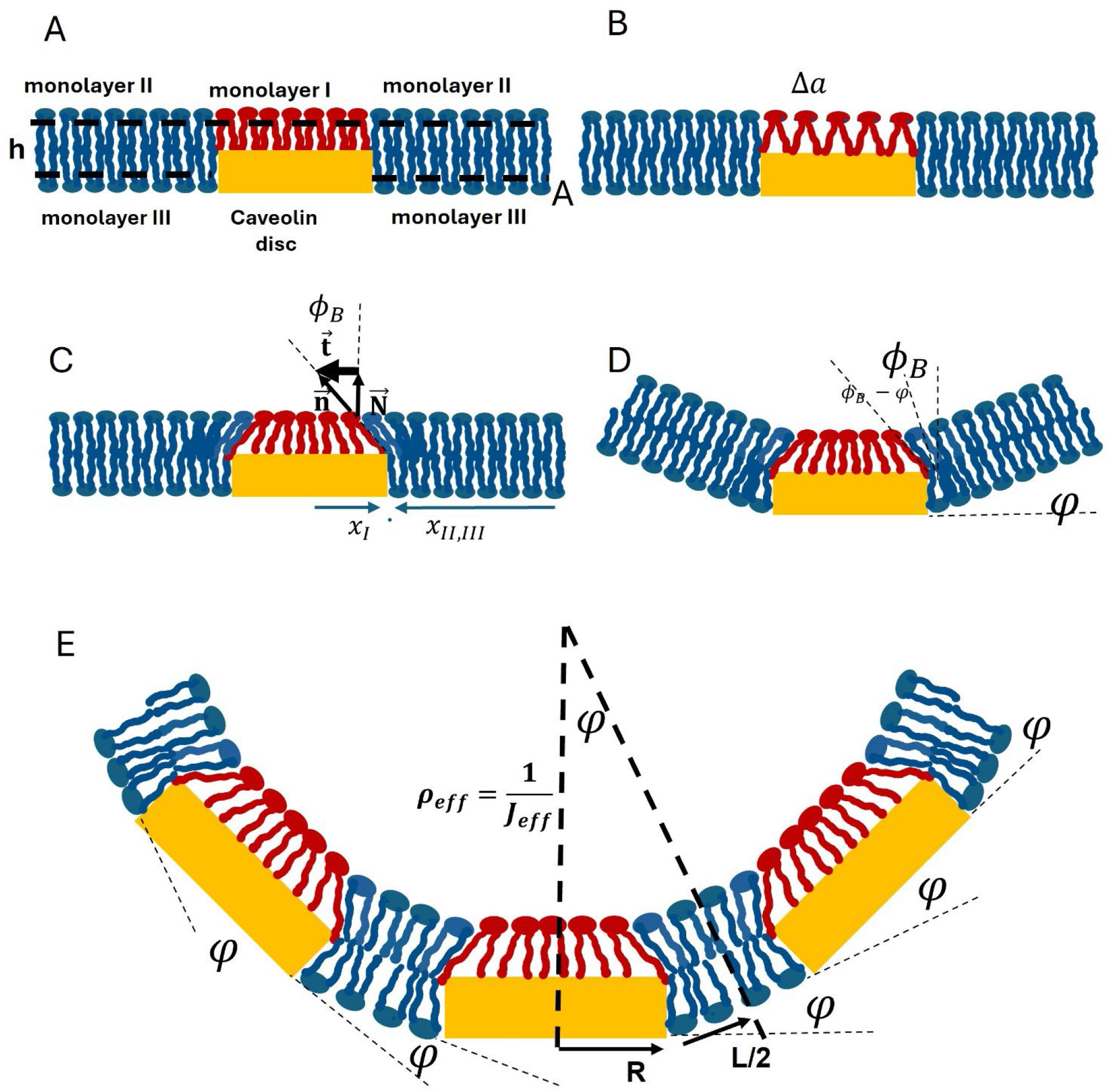
Description of the model. Yellow rectangles represent caveolin discs. The part/s of the distal monolayer contacting the hydrophobic plane/s of the disc/s (monolayer I) is/are shown in red colour. The monolayer fragment/s that do not contact the disc plane/s (monolayers II and III) is/are shown in dark blue colour. (A) The reference state preceding the relaxation of the system. The dashed lines represent the monolayer neutral planes. (B) Decrease of the number of lipid molecules contacting the disc that leads to the area stretching of the monolayer I. (C) Relaxation of the monolayer I stretching that leads to tilt and splay in the monolayers I and II. The axes x_II_ and x_II,III_ are chosen to describe, respectively, the monolayer I and the monolayers II and III; (D) Kinking of the membrane profile, which generates the tilt and splay in the monolayer III but results in an overall relaxation of the system’s elastic energy. (E) Membrane with multiple discs. Notations: ϕ_B_ and φin (C-E) are, respectively, the boundary tilt and kink angles; R and L/2 in (E) represent, respectively, the disc radius and the half-distance between adjacent discs determined along the system’s mid plane; the dashed lines in (E) show the boundaries of the building block defined as the system and at the same time represent the radius of the effective curvature, 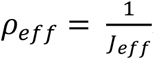.

The activity of caveolins and the accessory proteins in caveola biogenesis is significantly affected and modulated by the membrane lipids with a special role played by cholesterol ^6,16^. As shown by lipidomic studies of purified caveolae, the lipid composition of caveolar membranes is significantly different from that of the surrounding plasma membrane and characterised by elevated molar ratios of cholesterol, sphingomyelin, glycosphingolipids and gangliosides ^17-19^, with an estimated number of cholesterol molecules per caveola as large as 22,000 ^17^. In addition, the cytoplasmic caveolar leaflet may be enriched in phosphatidylserine^20^ and phosphoinositide (PI(4,5)P_2_) ^21^. Cholesterol is required to assemble 8S caveolin complexes into 70S complexes and stabilise the latter ^22^. Depletion of cholesterol from the plasma membrane leads to caveolar instability and flattening of the caveolar invaginations ^23,24^. Moreover, cholesterol addition to cells was shown to cause increased caveola curvature and caveolar budding ^19^. The mechanisms of recruitment of cholesterol and the specific phospholipids to caveolar membranes are poorly understood. Cholesterol can directly interact with caveolins ^18,25,26^, while cavins are able to bind phosphatidylserine and PI(4,5)P_2_ ^27,28^. Besides these apparently specific interactions, the partitioning of certain lipids to caveolae may be driven by effective interactions of non-chemical character resulting from perturbations of the membrane lipid matrix by the embedded caveolin complexes ^6^.

Caveolae generated by caveolins, with or without cavins, have peculiar configurations. While, initially, the shapes of caveolar bulbs were regarded as smoothly rounded, high-resolution EM imaging revealed the bulb profiles to have a faceted appearance ^10,29,30^. The straightforward intuition (but also see ^31^) suggests that the faceted shapes of caveolae are a direct consequence of the flatness of the caveolin discs ^13,29^. According to this logic, the caveolar facets must be paved by the discs ^13,14^, whereas the membrane gaps between the flat facets must be kinked to generate the overall closed membrane configuration.

The outlined experimental information and qualitative ideas on the caveolin discs driving caveola formation, the faceted appearance of caveolar bulbs, and the strong effect of cholesterol on the caveola stability pose a challenge to understanding the physical mechanism behind all this phenomenological knowledge. The outstanding specific issues are about the forces driving the membrane kinking between the caveolin discs necessary for the generation of the overall curved shapes of the caveolar surfaces, and the mechanism by which lipids, and particularly cholesterol, influence caveolar formation and disassembly.

Here we consider a model addressing these questions and provide a physics background for understanding the caveola shaping by caveolins. The central hypothesis of our model is that the interaction energy of the extracellular membrane leaflet with the hydrophobic plane of a caveolin disc differs from that with the cytoplasmic leaflet. We show that the difference in these energies drives the generation of elastic stresses and strains within the membrane monolayers, which, in turn, favour the membrane kinking at the disc boundaries and, hence, drive the overall curvature of caveolar membranes. We predict that these elastic stresses drive enrichment of cholesterol in the extracellular leaflets of caveolar membranes. We demonstrate that cholesterol facilitates the concentration of the caveolin discs in the membrane plane into dense domains and enhances caveolar membrane curvature. We demonstrate that cholesterol facilitates the concentration of the caveolin discs in the membrane plane into dense domains and enhances caveolar membrane curvature. The model does not rely on any specific interaction between cholesterol and other membrane components but rather uses only the cholesterol’s structural feature of having a strongly negative intrinsic molecular curvature^32^. Therefore, we predict that any other lipid of similar intrinsic curvature, such as DAG^33^, can have the same effect as that of cholesterol on caveola biogenesis.

### Qualitative essence of the model

To qualitatively explain the central idea of the model, we start by considering the effect of embedding a single disc into one of the membrane monolayers of an initially flat lipid bilayer. We discuss the factors leading to the kinking of the membrane profile along the edge of the disc, which can be seen as the generation of an overall membrane curvature. Further, we qualitatively consider the curved membrane configurations resulting from the insertion of multiple discs into the same monolayer and discuss the effective interaction between the discs expected to be mediated by the monolayer elastic stresses. We qualitatively discuss the expected impact of cholesterol (or other lipids with similar intrinsic molecular curvature) on the membrane shaping by the inserted caveolin discs.

#### Membrane with a single caveolin disc

The membrane monolayer harboring the disc will be referred to below as the proximal monolayer whereas the second monolayer will be called the distal one. We assume that the hydrophobic plane of the disc inserted into the proximal monolayer reaches the interface between the monolayers and, therefore, contacts the hydrophobic plane of the distal monolayer (Fig.1A). The fragment of the distal monolayer contacting the disc will be referred to as the monolayer I. The rest of the distal monolayer will be called the monolayer II. The part of the proximal monolayer flanking the disc will be referred to as the monolayer III (Fig.1A).

Our major assumption is that the contact between the distal and the proximal monolayers along the bilayer mid-plane is energetically more favorable than the contact between the distal monolayer and the hydrophobic plane of a caveolin disc. We will denote the former and latter contact energies related to one lipid molecule by, respectively, *ε*_*L*_ and *ε*_*P*_. The origin and driving force of all the effects considered below is the contact energy difference,

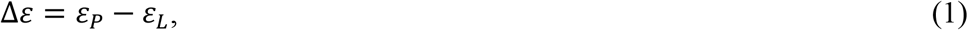

which is assumed to be positive, Δ*ε* > 0. In the language of physical chemistry, the contact energy is equivalent to wetting energy, so we assume that the wetting by the distal monolayer’s lipids of the hydrophobic surface of the proximal monolayer is preferable to wetting of the surface of the caveolin disc with Δ*ε* equivalent to the difference between the wetting energies per molecule.

In the following, we will refer to Δ*ε* as simply the contact energy, for brevity.

The chief idea of our model is that the differential wetting quantified by the contact energy, Δ*ε*, causes the generation of elastic deformations and stresses within the membrane monolayers, which, in turn, drive the kinking of the membrane profile at the disc’s edge. To explain the model intuitively, we decompose the kinking process into sequential steps.

In the first step, the caveolin disc of the area, *A*_*p*_, is inserted into the initially flat membrane but the system is kept flat, and the distal monolayer remains intact (Fig.1A). We denote by *N*_0_ the number of lipid molecules found at this stage in the monolayer I so that the in-plane area per lipid molecule, *a*, in this monolayer is 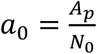. Here and in the following the area per lipid molecule, *a*, is determined at a special intra-monolayer plane referred to as the neutral plane for which the deformations of stretching are energetically decoupled from the deformations of bending ^34,35^ and, hence, of deformation of splay of the lipid hydrocarbon chains^36,37^(see below). The neutral plane lies, approximately, along the interface between the polar and hydrophobic moieties of a lipid monolayer (Fig.1A)^33,34,38^.

At the second step, because of the positive contact energy, Δ*ε* > 0 (Eq.1), the system tends to reduce the number of lipid molecules in the monolayer I. This can be achieved by shifting some number of lipid molecules, Δ*N*, from the monolayer I into the monolayer II (Fig.1 B). As a result of such shifting, the hydrophobic plane of the disc, *A*_*p*_, turns to be contacted by a smaller number, *N*_0_ − Δ*N*, of lipid molecules meaning that the monolayer I undergoes stretching (Fig.1B). The in-plane area per one lipid molecule in the monolayer I increase and becomes, *a* = *a*_0_ + Δ*a*, with the area increment approximately equal 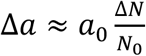. The monolayer stretching is associated with an elastic energy^39,40^, which is determined by the extent of deformation, 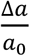, and the monolayer rigidity for stretching characterized by the stretching modulus, Γ ^39,40^.

The stretching energy accumulated at the second step can be partially or almost fully released at the third step in which the lipid molecules located at the boundary between the monolayers I and II (i.e., along the edge line of the caveolin disc) tilt towards the center of the disc (Fig.1C). This tilting enables the neutral plane of the monolayer I to contract and, hence, the lipid stretching, Δ*a*, to relax while keeping the same reduced number of lipid molecules, *N*_0_ − Δ*N*, in the monolayer I. We will quantify the extent of this tilt by the angle, *ϕ*_*B*_, referred to below as the boundary tilt (Fig.1C).

The larger *ϕ*_*B*_ the more efficient the relaxation of the monolayer stretching and the reduction of the stretching energy. Yet, lipid tilting requires an alternative energy cost. Being induced at the boundary between the monolayers I and II the lipid tilt propagates ^41,42^ into each of these monolayers, its value decreasing with the distance from the boundary (Fig.1C). The tilt variation along the monolayer surface gives rise to the splay deformation, whose essence is the relative spreading of the lipid polar heads and the ends of the hydrocarbon chains^36,37,42^ (Fig.1C). The splay is conventionally defined as positive if the lipid heads are spread compared to the chain ends and as negative otherwise^36,37^. The monolayer tilt and splay are associated with elastic energies, which are determined by the extent of each deformation and the related rigidity characterized by the elastic modulus *k*_*t*_ for tilt and *k* for splay ^36,37,42,43^.

Generally, the outcome of the second and third steps is a combination of stretching of the monolayer I and the tilt-splay in the monolayers I and II. Yet, the relative extent of these deformations is set by the relationships between their rigidities. The dominant deformation is the one for which the membrane monolayer has the least rigidity. Lipid monolayers are considerably less rigid for the tilt-splay deformation than for the area stretching. Indeed, the monolayer stretching modulus, Γ, is of the order of a few hundred 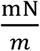 ^39,40^, whereas the tilt modulus, *k*, is in the range of a few tens 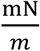 ^37,42-44^, and the splay modulus, *k* ≈ 4 10^−20^Joule ^45-47^related to the square of the monolayer thickness *h* ≈ 2nm is of the order of 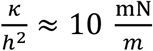. Therefore, the leading deformation resulting from the second and third steps must be that of the tilt-splay, while the monolayer stretching in the monolayer I must be negligibly small. In other words, the tilt-splay deformation at the third step must result in, practically, complete relaxation of the area stretching so that the latter will be neglected.

At the fourth step, we allow the system to deviate from the flat state by developing a kink of the membrane midplane ar the disc’s edge (Fig.1D) in case this can relax the tilt-splay energy. We quantify the extent of the kink by the kink angle, *φ*(Fig.1D). A simple geometrical consideration shows that for the monolayer II, the kinking reduces its tilt at the boundary with the monolayer I from *ϕ*_*B*_ to *ϕ*_*B*_ − *φ*(Fig.1D).

At the same time, the kinking results in the generation of the tilt-splay deformation in the monolayer III with the tilt at the boundary with the disc equal to the kink angle, *φ*. As a result, the kinking of the membrane profile reduces the elastic energy accumulated in monolayer II but generates some elastic energy in monolayer III. Yet, as we show by the calculations (see SI), the kinking enables a reduction of the overall elastic energy of the system and is, therefore energetically favorable.

Taken together, the major idea of the model is that the contact energy, Δ*ε* (Eq.1), favors a kinking of the membrane profile at the edge of the inserted caveolin disc (Fig.1D).

In the following sections, we quantitatively analyze the dependence of the kink angle, *φ*, on the contact energy, Δ*ε*, by using the models of the membrane tilt and splay elasticity.

#### Membrane with multiple caveolin discs

To grasp the qualitative essence of the membrane shape generated by multiple caveolin discs, we can consider it to be composed of elementary units, each unit represented by a bilayer fragment containing a single disc flanked by lipid bilayer fragments and having a kinked profile as discussed above (Fig.1D). Putting such units side-by-side as mosaic elements with a smooth transition between their stress and shape profiles provides an overall curved configuration (Fig.1E). The effective curvature of the membrane, *J*_*eff*_, is set by the kink angle, *φ*, and the effective arc length, which is equal to the sum of the disc radius and the half-distance between the discs (Fig.1E).

There must be a cross-talk between adjacent discs mediated by an interplay between the elastic stresses generated in the membrane matrix by each of them. This interplay is expected to affect the values of the kink, *φ*, and boundary tilt, *ϕ*_*B*_, angles at every disc making them dependent on the inter-disc distance, *L*. Moreover, the elastic stress interplay must mediate an effective interaction between the discs.

In the following sections, we will derive the quantitative relationships relating the geometrical parameters of the system and the effective interaction between the discs to the inter-disc distance by using the model for the tilt and splay elasticity of lipid monolayers^36,37,42^.

#### Effects of cholesterol

In the above reasoning, we implicitly assumed the membrane monolayers to be homogeneous all over the monolayer areas. This consideration did not account for a possible interplay between the monolayer stresses induced by the insertion of the caveolin discs and a, generally, inhomogeneous distribution along the monolayer planes of lipid species modulating the monolayer elastic properties. For mammalian caveolae, the most essential lipid of this kind is cholesterol, which, as already mentioned, has a remarkable impact on the caveola morphology and stability ^17,19,22-24^. As far as the lipid monolayer elasticity is concerned a distinctive feature of cholesterol is its effective molecular intrinsic curvature, *ζ*, which is exhibited by cholesterol in mixed monolayers and has a strongly negative value in the range between 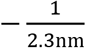 and 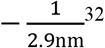. Due to this molecular feature, the presence of cholesterol in the lipid monolayer regions subjected to negative lipid splay decreases the related elastic energy^36,37^, this effect is proportional to the cholesterol surface concentration^48^. In the monolayer I of our system the hydrocarbon chains of lipid molecules are spread compared to the polar heads so that the splay is negative. In monolayers II and III the heads are spread relatively to the chain ends and, hence, the splay is positive (Fig.1C-E). Hence, an enrichment of the monolayer I and depletion of the monolayers II and III in cholesterol are expected to reduce the monolayer elastic energy and enhance the membrane kinking and tilting driven by the difference in the contact energies, Δ*ε*.

In the following, we quantitatively account for the interplay between the membrane stresses and the cholesterol surface concentration resulting in the cholesterol partitioning between the plasma membrane (that plays the role of a lipid reservoir) and different monolayers of the system. We analyze the impact of the cholesterol partitioning on the extent of the overall membrane curvature and the effective interaction between the embedded caveolin discs.

Importantly, as already mentioned, the results we obtained for cholesterol must be valid for any other lipid with a strongly negative intrinsic curvature. For example in sterol-free *E*.*coli* ^10^ this requirement may be provided by diacylglycerol (DAG) ^33^.

### Main definitions and equations

#### Description of the system

We consider a lipid bilayer with caveolin discs of radius *R* inserted into its proximal monolayer and separated by a distance *L* along the bilayer midplane. We assume, for simplicity, that the bilayer consists of two types of lipids, the basic lipid and cholesterol. The local molar ratio of cholesterol in a monolayer will be denoted by *c* so that the molar ratio of the basic lipid is 1 − *c*. Each monolayer of the bilayer can freely exchange the lipid material with an external reservoir consisting of the same components. The molar ratio of cholesterol in the reservoir is *c*_0_. The reservoir is a prototype of the plasma membrane to which real caveolae are connected.

For technical simplicity, we consider the two-dimensional version of the model in which a bilayer’s configuration is described by the two-dimensional profile of its mid-surface (Fig.1) and is uniform in the third direction. We expect the model predictions to retain their qualitative validity also for realistic three-dimensional configurations.

The bilayer with the inserted discs can be considered as built up of structural blocks (Fig.1E). Each such block consists of half of a caveolin disc of length *R* covered by the monolayer I, and the bilayer fragment of length *L*/2, consisting of the monolayers II and III and extending halfway to the adjacent disc (Fig.1E). The structural block will be referred to below as the system, for brevity. The point on the system’s midplane separating the disc and bilayer parts (Fig.1E) will be called the internal boundary of the system. The external boundaries of the system are the points on the mid-plane representing the interface between the system and the adjacent structural blocks.

In the following, the physical values of the monolayers I, II, and III of the system will be denoted by the corresponding index. The relationships in which no indices are used are applicable to any of the three monolayers.

Each monolayer of the system is described by its neutral plane, which is parallel to the system’s midplane and separated from it by a distance, *h*, approximately equal to the monolayer thickness (Fig.1A). Each is characterized at every point of its neutral plane by the tilt vector, 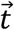 and splay, 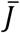, of the constituent lipid molecules. A detailed introduction of the notions of tilt and splay is presented and discussed in ^36,37,41,42^. In brief, the lipid tilt, 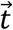, quantifies the deviation of the unit vector, 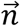, describing the orientation of the lipid hydrocarbon chains from the unit normal, 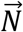, of the monolayer neutral monolayer plane (Fig.1C). For the 2D model considered here we choose the x-axis directed along the neutral plane from the external to the internal boundary. For small angles, *ϕ*, between 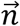 and 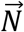, the projection of the tilt vector to the x-axis is equal *t* = *ϕ* if 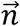 slants away from 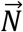 in the direction of the x-axis (Fig.1C) and, *t* = −*ϕ*, for the opposite direction of the slanting. The lipid splay,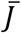, quantifies the variation of the lipid chain orientation along the monolayer neutral plane and is determined as the sum of the divergence of the tilt, 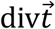, and the mean curvature of the monolayer neutral plane, *J* ^36,37,41,42^. As we will explain below, in the 2D model considered here, the curvature can be neglected so that all three monolayers of the system will be regarded as flat, *J* = 0, in which case the splay is equal to, 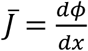, with *x* being the coordinate along the monolayer x-axis.

The physical state of each of the system’s monolayers is characterized by the distribution along the monolayer neutral plane of the lipid tilt angle, *ϕ*(*x*), the splay, 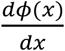, and the cholesterol concentration, *c*(*x*).

The system’s mid plane profile sharply turns at the internal boundary. This turn will be described by the kink angle, *φ*(Fig.1D,E). In addition, we define the boundary tilt angle, *ϕ*_*B*_, as illustrated in (Fig.1D,E). The values of *ϕ*_*B*_ and *φ*set the tilt angles in the three monolayers at the internal boundary of the system,

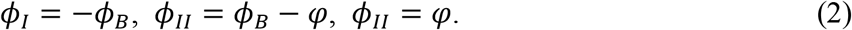

At the external boundaries of the system, all the tilt angles must vanish,

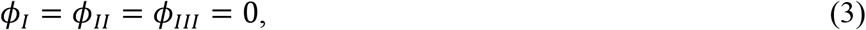

to guarantee a smooth transition to the adjacent structural blocks.

#### Energy of the system

Our goal is to find the equilibrium configuration of the system corresponding to the minimum of the system’s energy, *F*. The energy will be computed with respect to the initial state in which the system is flat so that the kink angle vanishes, *φ*= 0, and there is no tilt-splay all along each of the three monolayers which also means that the boundary tilt angle vanishes *ϕ*_*B*_ = 0 (Fig.1A). The cholesterol concentration, *c*, i n the initial state is homogeneous in each monolayer and equal to that in the reservoir, *c*_0_.

We consider the energy *F* to consist of the change of the contact energy, *F*_*cont*_, the monolayer elastic energies of the emerging tilt and splay, *F*_*el*_, and the energy of cholesterol repartitioning between the system’s monolayers and the reservoir, *F*_*chol*_, so that

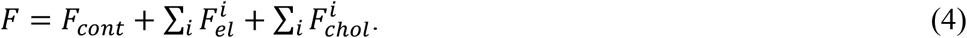

Here and below, the index “i” becomes I, II, or III depending on system’s monolayer in mind. The meaning of Σ_*i*_ in (Eq.4) is the summation of the contributions by the three monolayers of the system.

The change in the contact energy, *F*_*cont*_, is proportional to the variation in the number of lipid molecules contacting the hydrophobic planes of the caveolin discs (monolayer I), Δ*N*, which results from the emerging tilt deformation. For small boundary tilt angles, *ϕ*_*B*_<1, it can be expressed as

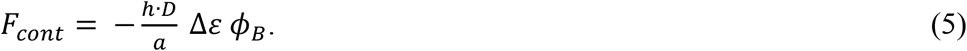

In (Eq.5) and below the multiplier *D* is used to convert the energies provided by the two-dimensional model and, thus, related to unit length of the axis pointing in the third direction, into the energies related to a caveolin disc and having the regular units. Therefore, the value of *D* will be taken equal to the size of a caveolin disc, *D* = 2*R*. The area per lipid molecule on the monolayer neutral plane, *a*, and the monolayer thickness, *h*, will be assumed constant because of the effective non-stretchability of the monolayer.

To determine the energy of tilt and splay in each monolayer of the system we will use the elastic model introduced and developed in^36,37^ and recently summarized in detail in the Supplementary Material of ^41^. This energy is related to the unit area of the monolayer neutral plane for small deformations, *ϕ*<1 and 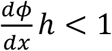,is given by

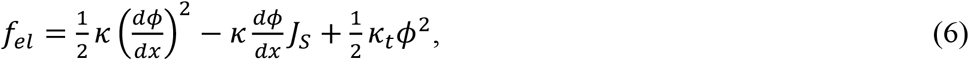

where *κ* is the monolayer splay modulus equivalent to the Helfrich bending modulus, *κ*_*t*_ is the monolayer tilt modulus, and *J*_*S*_ is the spontaneous splay equivalent to the Helfrich spontaneous curvature^36,37^. The spontaneous splay, *J*_*S*_, depends on the intrinsic curvatures of the constituent lipids and the lipid composition ^49^. We use the simplest linear model for this dependence^49,50^ according to which,

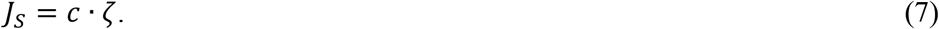

where *ζ* is the intrinsic curvature of cholesterol. The relationship (Eq.7) implies that the basic lipid has a vanishing intrinsic curvature. The total elastic energy of a monolayer is obtained by the integration of *f*_*el*_ over the monolayer area, *A*, which for the 2D model is reduced to the integration over the linear coordinate, *x*,

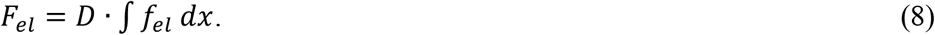

The energy of cholesterol redistribution, *F*_*chol*_, is the consequence of a decrease in the entropy of mixing between cholesterol and the basic lipid, which accompanies a deviation of the mole fraction of cholesterol, *c*, from that of the reservoir, *c*_0_. The resulting change of the free energy related to the unit area of the monolayer neutral plane can be presented as (see e.g. SI of ^51^ for derivation and detailed explanations),

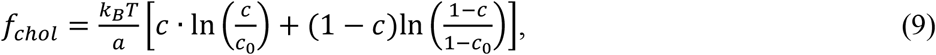

where *k*_*B*_*T* is the product of the Boltzmann constant and the absolute temperature, and *c* is the local mole fraction of cholesterol varying along the monolayer’s neutral plane. The relationship (Eq.9) assumes that the molecular in-plane area, *a*, of cholesterol is equal to that of the basic lipid, which is a simplifying approximation.

We assume that the deviation of the cholesterol molar ratio, *c*, from its reservoir value, *c*_0_, is small, 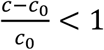, all over the system and use the first non-vanishing order approximation of (Eq.9),

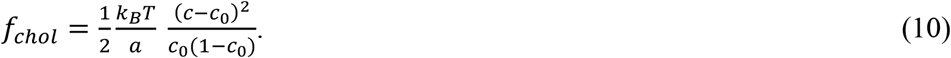

The total energy of cholesterol repartitioning is given by the integration of *f*_*chcee*_ (Eq.10) over the area of the monolayer neutral plane,

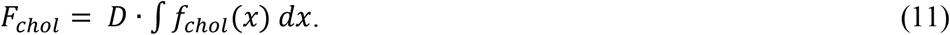

#### Strategy of analysis

We will start with determining the equilibrium distributions of the tilt angle, *ϕ*(*x*), the splay, 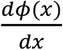, and the cholesterol concentration, *c*(*x*), along the neutral plane of each monolayer for fixed values of the kink angle, *φ*, and the boundary tilt angle, *ϕ*_*B*_, and a given half-distance between the adjacent discs, *L*/2. This will be done by deriving the equations of equilibrium for the tilt and cholesterol distributions and solving them for each monolayer upon fulfilling the boundary conditions (Eqs.2,3) (see SI A). Using the obtained results and (Eqs.4-11), we will determine the energy of the system as a function of the fixed values, *F*(*ϕ*_*B*_, *φ, L*/2). Further, for each distance, *L*, between the caveolin discs, we will seek the values of the kink, *φ*, and boundary tilt, *ϕ*_*B*_, angles minimizing the energy.

## Results

For each inter-disc distance, *L*, we describe the equilibrium configuration of the system by the optimal value of the kink angle, *φ*^∗^ (Fig.1D,E), determining the system’s shape, the optimal value of the boundary tilt angle, 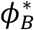 (Fig.1D), characterizing the extent of the tilt and splay deformations in the system, and the optimal deviations of the distribution of the cholesterol mole fractions, *c*, in the system’s monolayers. In addition, we determine the membrane-mediated force acting between the discs, which will allow us to predict whether the assembly of caveolin discs into caveolar coats can occur spontaneously or requires the action of additional forces counteracting the membrane-mediated ones. The full details of the computations are presented in (SI A). Below we present only the results of this analysis.

### Possible instability with respect to kinking

The first important conclusion of our analysis (SIA) is that the cholesterol redistribution causes an effective softening of the system’s monolayers for the splay deformations. As a result, the system’s resistance to the splay and the related elastic energy are determined by an effective splay modulus, *κ*_*eff*_, which is expressed through the basic splay modulus, *κ*, appearing in (Eq.6), the cholesterol mole fraction in the reservoir, *c*_0_, and the molecular intrinsic curvature of cholesterol, *ζ*, by

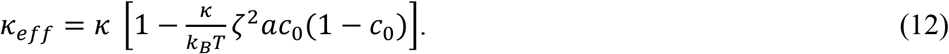

According to (Eq.12), the effective splay modulus, *κ*_*eff*_, decreases with the increase of the cholesterol mole fraction, *c*_0_, within the biologically relevant range, 0<*c*_0_<0.5. Importantly, for certain values of the cholesterol mole fraction, *c*_0_, the effective splay modulus can adopt vanishing or negative value, *κ*_*eff*_ ≤ 0. In this case, the system is predicted to tend to develop unlimitedly large boundary tilt, *ϕ*_*B*_, and kink, *φ*, angles. The possibility of this instability is controlled by *c*_0_ and the parameter combination,

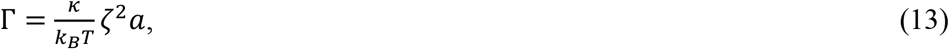

the latter having a meaning of the ratio between the characteristic elastic energy, *kζ*^2^*a*, of deforming one cholesterol molecule from the stress-free state of the intrinsic splay *ζ* to the state of vanishing splay, and the thermal energy, *k*_*B*_*T*.

The most favorable for the instability is the cholesterol mole fraction 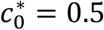. According to (Eq.12), at this level of cholesterol the onset of instability is possible if the parameter Γ is equal or exceeds a critical value Γ^∗^,

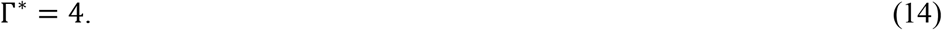

For lower cholesterol content, *c*_0_<0.5, the instability requires even larger values of Γ > Γ^∗^.

The value of the intrinsic curvature of cholesterol, *ζ*, was measured to be between 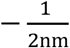 and 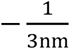^32^. Using for the cholesterol molecular area the value *a* = 0.7 *nm*^−2^, the criterion of instability (Eq.19) is fulfilled if the monolayer splay modulus *κ* is larger than a threshold value that varies in the range between 20 - 50 *k*_*B*_*T* depending on *ζ*. This may be possible for monolayers containing large amounts of charged lipids ^52^ and/or perhaps of cholesterol^53^. For more common values of the monolayer splay modulus, *κ*, varying in the range between 10 – 15 *k*_*B*_*T*, the parameter Γ is smaller than the critical value, Γ<4, so that the system must be stable for all cholesterol mole fractions.

In the following, we use Γ = 0.35 which, on one hand, guarantees the stability of the system in the whole feasible range of the cholesterol mole fraction, *c*_0_<0.5.

### Characteristic scales

To simplify the understanding of the following results (SI A), it is convenient to define the combinations of the system’s parameters which set the energy and length scales of the system. We classify the scales into the basic and operational ones.

The basic scales are determined by the basic elastic parameters introduced in the previous section. The basic energy scale, σ_0_, is set by the basic monolayer splay, *κ*, and tilt, *κ*_*t*_, moduli along with the monolayer thickness, *h*, and the in-plane area per lipid molecule, *a*,

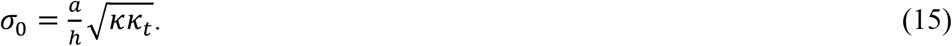

The reported values of the monolayer splay (bending) modulus, *κ*, determined for monolayers of different lipid compositions ^47,54^, various membrane charges^52^, and diverse cholesterol contents^53,55,56^ vary in a wide range between less than 10*k*_*B*_*T* and about 30*k*_*B*_*T*. The reported values of the tilt modulus, *κ*, are in the range 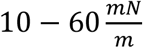^37,42-44^. Since *h* ≈ 2*nm*, and *a* ≈ 0.7*nm*^2^, the basic energy scale σ_0_ is the range about 2 – 20 *k*_*B*_*T*.

The basic length scale whose meaning is a characteristic decay length of the splay and tilt deformations is,

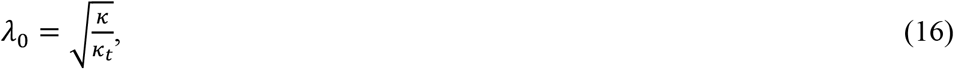

and has values within the approximate range of 1 to 3nm.

The operational energy, σ, and length, *λ*, scales account for the effects of the cholesterol repartitioning between the system’s monolayers and the reservoir. They are determined, respectively, by the relationships (Eq.15) and (Eq.16) in which the splay modulus, *κ*, must be replaced by its effective value (Eq.12),

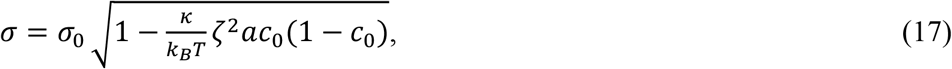

and

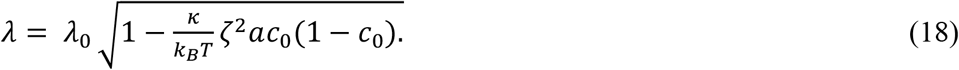

According to (Eqs.17,18) the cholesterol redistribution reduces both the characteristic energy of the system and the decay length of the elastic stresses.

### Optimal configurations

The optimal shape of the system for a given inter-disc distance, *L*, is set by the optimal kink angle, which is given by (see SIA)

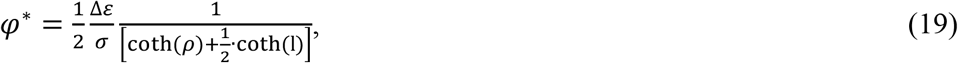

while the optimal boundary tilt angle

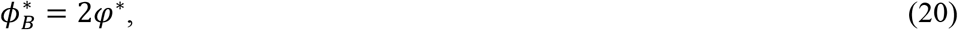

where σ is the operational energy scale (Eq.17), and *l* and *ρ* are, respectively, the half-distance between adjacent caveolin discs and the disc radius both normalized by the operational length scale (Eq.18)

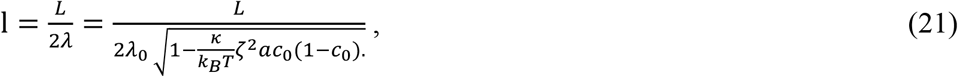

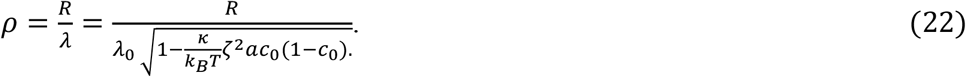

The dependence of the optimal kink angle, *φ*^∗^, on the inter-disc distance, *L*, for different values of the cholesterol mole fraction, *c*_0_, is presented in (Fig.2A).

**Figure 2.**
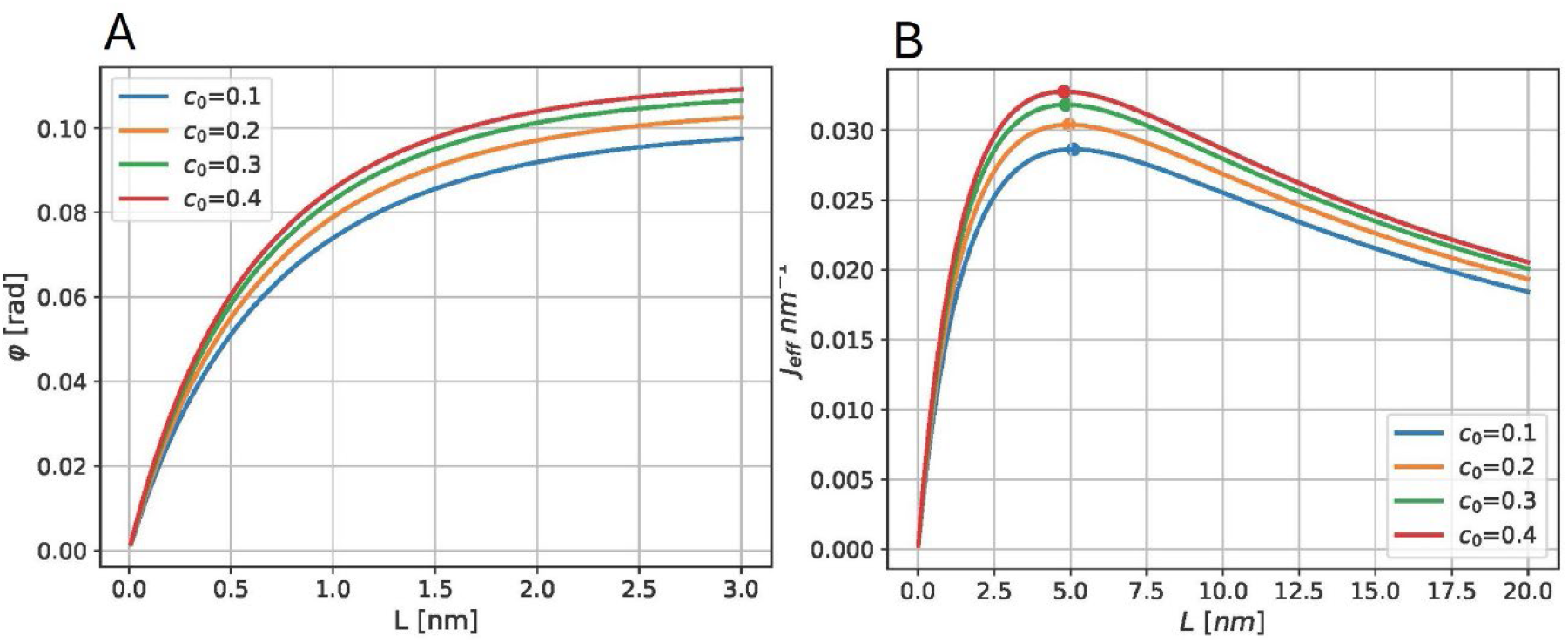
The dependences of the optimal geometrical parameters of the system on the inter-disc distance, L, for different values of cholesterol in the reservoir, c_0_. (A) The kink angle, φ; (B) The effective curvature, J_eff_. Dot at each curve indicates the distance of a maximal effective curvature. Parameters used for all curves: ζ = 1/2.5 nm^−1^, k = 10k_B_T, k_t_ = 10 k_B_T/nm^2^, Δε = 1k_B_T, R = 7nm, h = 2nm, a = 0.7nm^2^.

The resulting effective curvature of the faceted system’s profile can be determined as

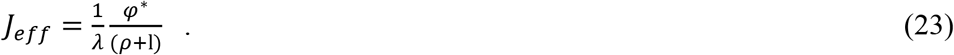

The dependence of *J*_*eff*_ on the inter-disc distance, *L*, for different *c*_0_ is presented in (Fig.2B).

These results (Fig.2) show that both the optimal kink angle, *φ*^∗^, and the corresponding effective curvature of the system, *J*_*eff*_, increase with rising cholesterol mole fraction, *c*_0_. This means that cholesterol is predicted to boost the membrane kinking and increase the effective membrane curvature generated by the caveolin discs.

A remarkable feature of our results is that the effective curvature, *J*_*eff*_, reaches a maximal value, 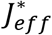, for a certain inter-disc distance, *L*^∗^, which, therefore, is the most favorable in terms of the curvature generation. The representative dependences of 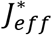 and *L*^∗^ on the cholesterol mole fraction *c*_0_, for typical values of the basic elastic parameters are presented in (Fig.3).

**Figure 3.**
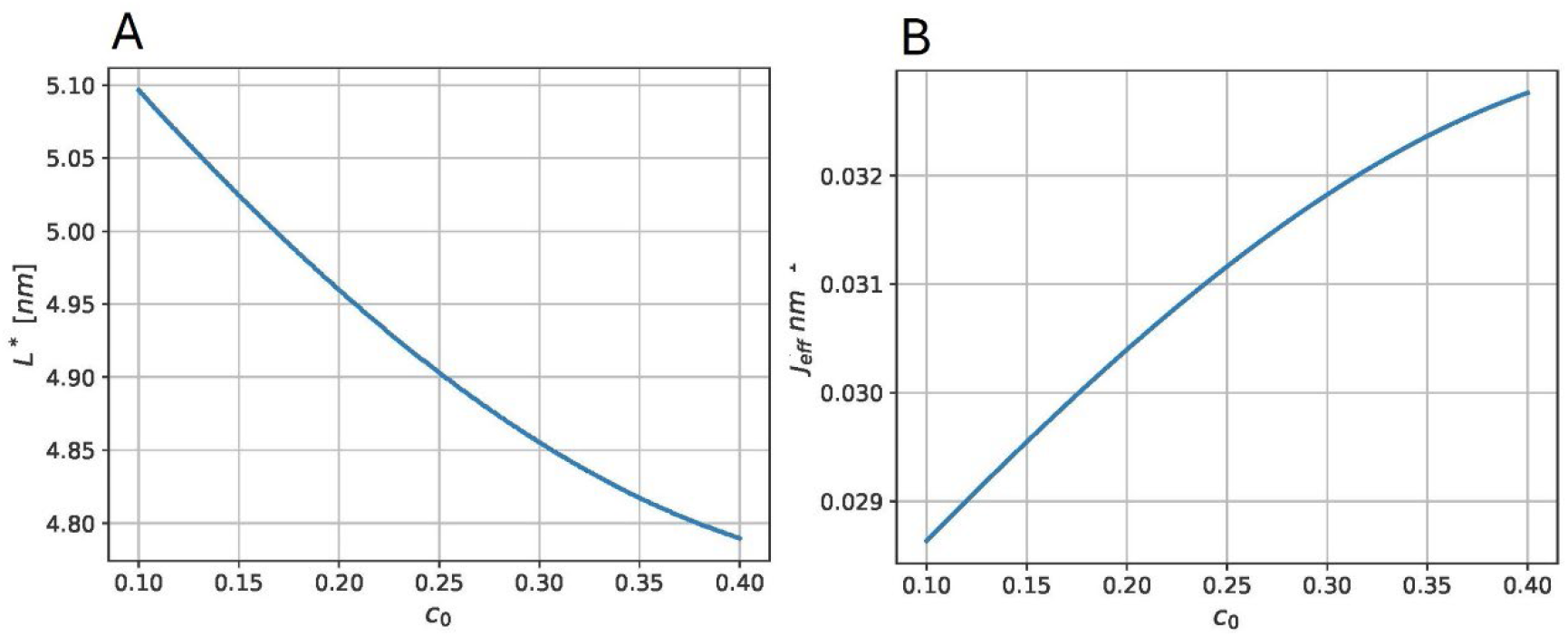
The optimal conditions for the curvature generation for different mole ratios of cholesterol, c_0_. (A) The optimal inter-disc distance, L^∗^, at which the effective curvature adopts a maximal value (B)The maximal effective curvature 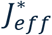. The parameter values used: ζ = 1/2.5 nm^−1^, k = 10k_B_T, k_t_ = 10 k_B_T/nm^2^, Δε = 1k_B_T, R = 7nm, h = 2nm, a = 0.7nm^2^.

Based on the observations of faceted caveolae ^10,29,30^, the distances between the flat facets, *L*, are in the range of a few nanometres while the effective caveolar radii are nearly 30nm. According to our model predictions (Figs.3A), such distances are optimal for curvature generation. Another conclusion of our results (Figs.3B) is that for a modest value of the contact energy Δ*ε* = 1*k*_*B*_*T*, the observed 30nm radii of caveolar membranes correspond to biologically feasible values of the cholesterol mole fraction, *c*_0_, in the range of 20-30%. In case of the contact energy, Δ*ε*, larger than 1*k*_*B*_*T*, lower cholesterol mole fractions would be required for generation of the observed curvatures.

The distributions of the cholesterol mole fraction, *c*, in the system’s monolayers (see SIA, Eqs. A12, A16, A20)) are illustrated in (Fig.4). The monolayer I contacting the hydrophobic face of the caveolin disc is enriched in while the monolayers II and III bridging the adjacent discs are depleted from cholesterol. The major enrichment and depletion are concentrated near the disc boundary.

**Figure 4.**
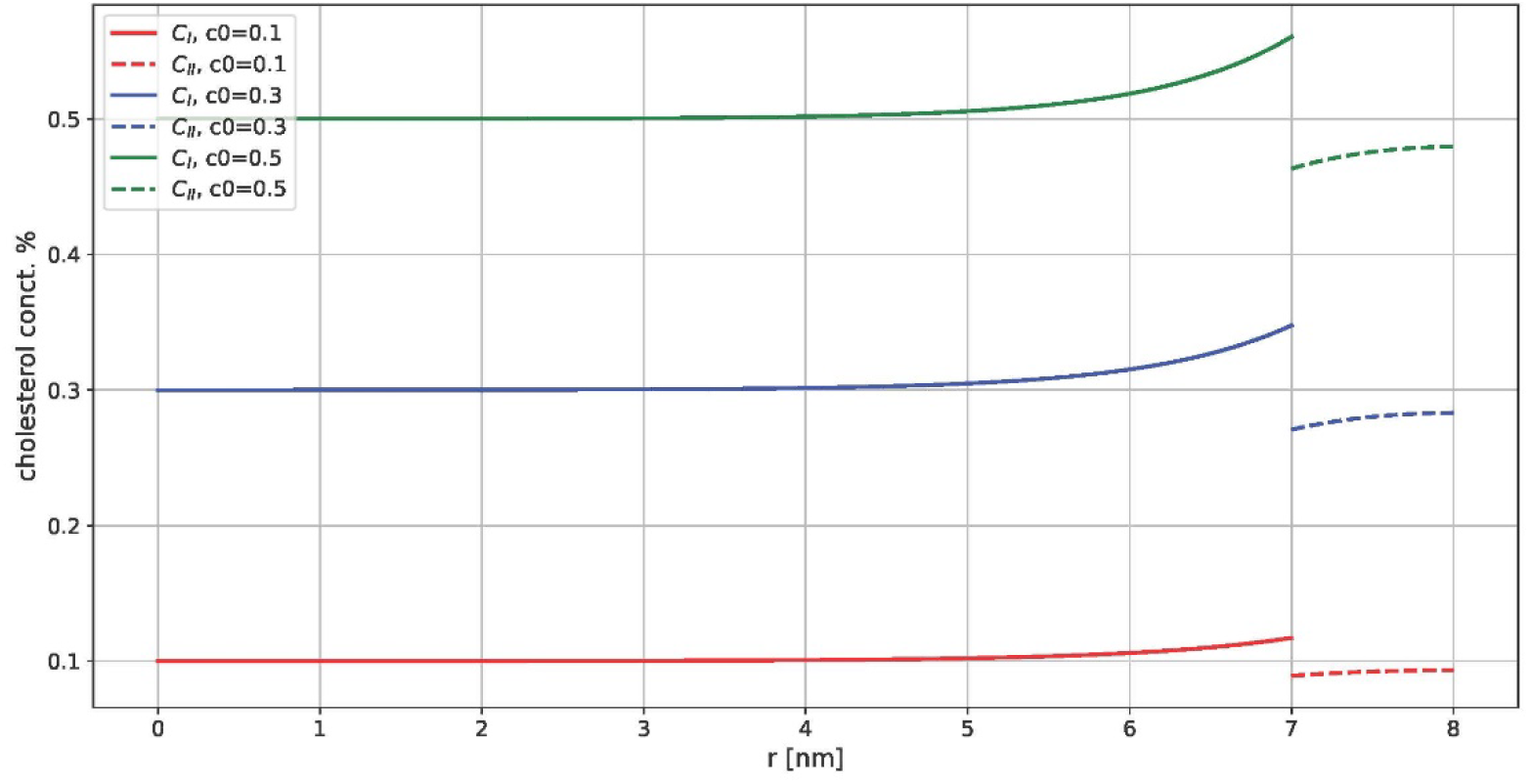
Distribution of cholesterol mole fraction across the system for different values of the background mole fraction, c_0_. The values of the boundary tilt ϕ_B_ and kink φangles correspond to the energy minimum. The solid lines describe the area of the monolayer II, the dashed lines describe the monolyers IIII and IIIIII. The parameter values used for all curves: k = 10k_B_T, k_t_ = 10k_B_T, ζ = 1/2.5nm^−1^, Δε = 1k_B_T, L = 2nm, R = 7nm, h = 2nm, a = 0.7nm^2^.

### Membrane-mediated interaction between the caveolin discs

The dependence of the system’s energy on the inter-disc distance is given by (see SI A)

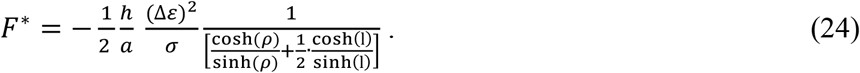

The membrane-mediated repulsive force acting between the adjacent caveolin discs, ℱ, corresponding to the energy (Eq.24) is given by

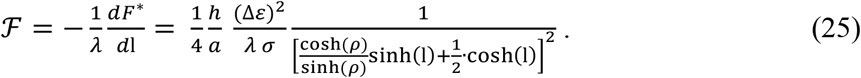

The dependences of the energy and force on the inter-disc distance, *L*, are presented in (Fig.5).

**Figure 5.**
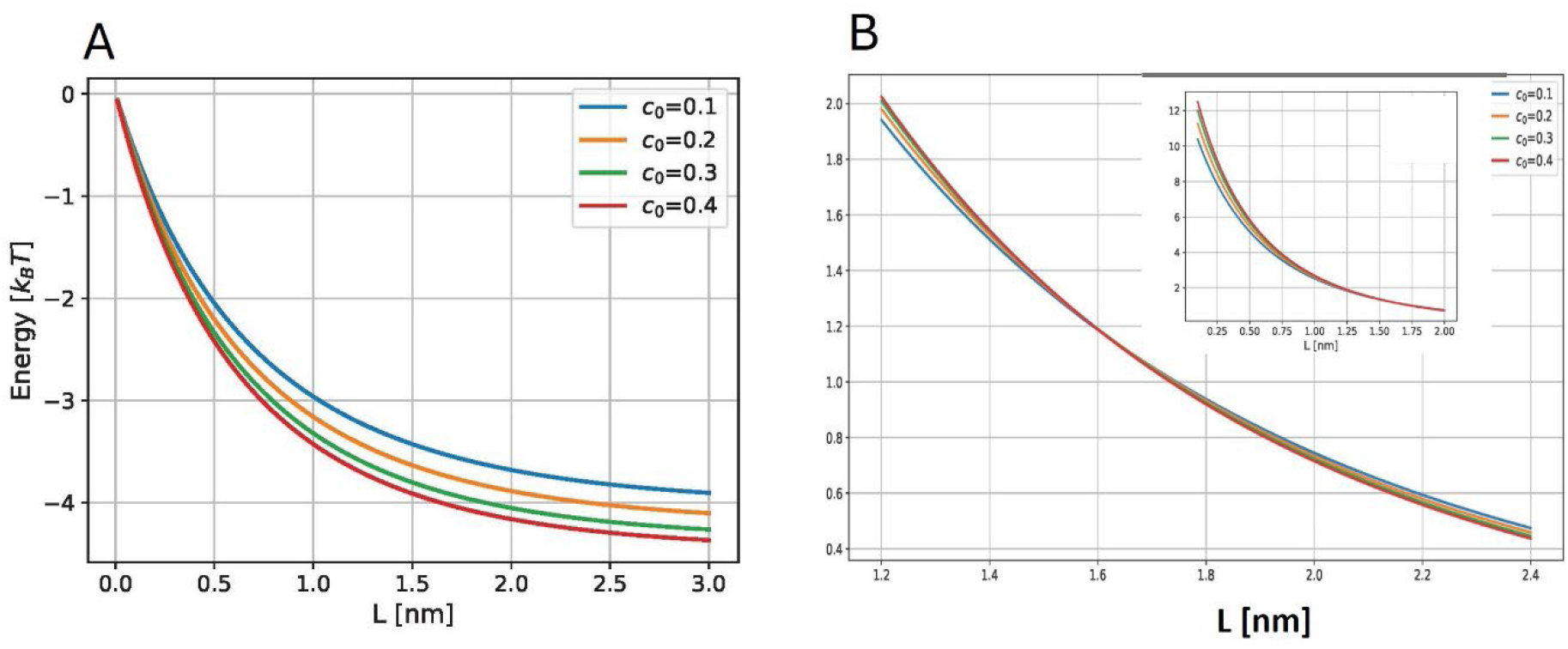
The membrane-mediated interaction between the caveolin discs. The energy of the system (A), and the corresponding repulsive force between adjacent discs (B) as functions of the inter-disc distance, L, for different values of cholesterol in the reservoir, c_0_. Parameters used for all curves: ζ = 1/2.5 nm^−1^, k = 10k_B_T, k_t_ = 10 k_B_T/nm^2^, Δε = 1k_B_T, R = 7nm, h = 2nm, a = 0.7nm^2^, D = 14nm.

The energy, *F*^∗^ (Fig.5A) is negative, which is a result of an overall relaxation of the energy of the initial state due to the emerging tilt, splay and kink deformations in the system’s monolayers, as qualitatively discussed above. The most efficient relaxation is reached for the inter-disc distances exceeding the operative length scale, *L* ≫ *λ*. Increase of the cholesterol mole fraction, *c*_0_, results in deeper energy relaxation due to a reduction of the energy cost of the splay deformation (Fig.5A).

The membrane-mediated force between the discs, ℱ, is repulsive for all inter-disc distances (Fig.5B). The predicted repulsive force is, practically, zero for large inter-disc distances and starts to grow when the distance becomes comparable to or smaller than the operational length scale, *L* ≤ *λ*. Since *λ* decreases with a growing mole fraction of cholesterol, *c*_0_ (Eq.18), a rise in the membrane cholesterol concentration is predicted to reduce the distance to which the discs can approach each other without any significant membrane-mediated resistance.

To get a sense of the characteristic magnitude of the membrane-mediated repulsive force between the caveolin discs, ℱ, we estimate its value for an experimentally relevant distance of say *L* = 2nm. According to (Fig.5B), the force is 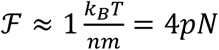. This is a biologically feasible force, which can be developed, for example, by just a couple of molecular motors^57^.

## Discussion

The mechanisms by which the caveolin-based protein complexes sculpt the 50-80nm large caveolar bulbs have remained elusive despite a long history of the field spanning many decades ^58^. While the major ideas behind the mechanisms of membrane shaping by proteins have been elaborated ^58-60^, the shapes of caveolae pose an additional challenge related to their faceted appearance ^10,13,29,30^ uncommon to other types of membrane buds. Whereas cavins can control the caveolar shaping either through their binding to caveolins within a two-layer structure ^9^ or due to the electrostatic interactions mediated by the cavin disordered domains ^61^, the primary role must be played by caveolins which have been shown to generate the caveola-like structure if expressed in otherwise caveola-free bacterial membranes lacking cavins ^10^. The previously proposed mechanism by which a caveolin molecule alone could produce local membrane curvature was based on a computationally discovered binding between the caveolin molecules shallowly embedded into one membrane leaflet and the molecules of cholesterol constituting a considerable part of the membrane lipid composition ^62,63^. Yet a shallow embedding of caveolin needed for the membrane curvature generation ^64^ does not comply with the experimental findings ^6,7^and an ensemble of such caveolin-cholesterol molecular complexes should be expected to generate smooth rather than faceted membrane profiles. Moreover, caveolin generates caveola-like membrane invaginations when expressed in cholesterol-free *E*.*coli* ^10^.

The recent structural data on caveolin assembly and arrangement within membranes made the phenomenon of membrane curving by caveolins even more puzzling and suggested that it is driven by a mechanism different from all those considered previously ^13^. Caveolin was discovered to oligomerize into flat discs of about 14nm diameter inserted into one membrane leaflet up to the interface with the second leaflet. It feels counter-intuitive that such large disc-like particles lying flat on the membrane plane can drive membrane curvature.

Here we proposed a mechanism to explain this phenomenon. The central hypothesis of the model is that the contact energy of lipid molecules of one membrane leaflet with the hydrophobic plane of a caveolin disc embedded into the second leaflet is larger than the energy of the contact between the two leaflets. We showed by modeling that such differential contact interaction results in the kinking of the membrane profile along the boundaries of the inserted caveolin discs, and, hence, to an effective membrane bending. Importantly, such bending-by-kinking mode results in a faceted rather than smoothly curve membrane profiles, in accord with experimental observations ^10,29,30^. Using the geometrical parameters of cell caveolae and the established values of the elastic moduli of lipid monolayers, we concluded that the differential contact energy per lipid molecule of the order of 1*k*_*B*_*T* is sufficient to explain the observations.

In addition, our model accounted for an enhancing effect of cholesterol and/or other lipids, such as DAG, with similar intrinsic curvature on the effective membrane curving by caveolin discs.

### Model predictions

An important prediction of our model is that the membrane stresses generated by the caveolin disc insertion result in a repulsive interaction between the discs. This means that an attractive force, ℱ_*att*_, which is not a part of the presented analysis, must act between the caveolin discs to overcome this membrane-mediated repulsion and concentrate the discs in the membrane plane into domains with an experimentally observed high density of caveolin. There are different possible origins of such attractive force. One is an effective interaction mediated by inter-disc connections through cavin molecules in vertebrate cells. Another is non-specific Vander-Waals interaction ^65^.

The model predicts that the induced effective membrane curvature, *J*_*eff*_, non-monotonously depends on the distance, *L*, between the adjacent cavin discs (Fig.2B). The curvature vanishes for zero, *L* = 0, and infinite, *L* →∞, separations, and reaches a maximal value, 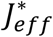, for the optimal distance, *L*^∗^, which is in the range of a few nanometers for a characteristic mole ratio of cholesterol, *c*_0_ = 0.3 and the typical values of the monolayer elastic parameters. This means that the most efficient curvature generation occurs if the attractive force, ℱ_*att*_, has such character and distance dependence that it brings the caveolin discs to a separation *L*^∗^ but not closer than that. The characteristic value of the attractive force needed to bring the caveolin discs to about 2nm separation is predicted to have a value of a few pN. In case of Vander-Waals interaction between discs, according to a rough estimation, such force corresponds to Hamaker constant *A*_*Ham*_ of a few tens of zeptojoules (zJ) which is feasible for interaction through a hydrophobic medium ^65^. If the attractive interaction between caveolin discs is provided by binding of the cavin proteins ^61^, relevant for the efficiency of the curvature generation is the length of a cavin-mediated bond between the discs determined by the detailed structure of the cavin-caveolin complex. This bond length must be in the nanometer range.

Essential predictions of the model relate to the effects and distribution of cholesterol. Generally, the model predicts a substantial enhancement by cholesterol of the caveolar membrane shaping by caveolin discs and provides a mechanistic background to all the observed effects of cholesterol on caveola formation described in the Introduction. Specifically, an increase of the background cholesterol mole fraction, *c*_0_, is predicted to boost the effective membrane curvature, *J*_*eff*_, generated by the discs (Fig.3B) which agrees with the observations ^19^. Further, an increase of *c*_0_ is predicted to reduce the membrane-mediated repulsive force between the discs at large and intermediate separations and, hence, to alleviate the requirement on the strength of an attractive force needed for a sufficient concentration of the caveolin disc in the membrane plane and generation of relevant membrane curvatures. This may explain why cholesterol was required for the assembly and stabilization of 70S caveolin complexes ^22^ as well as the observed caveolar instability and flattening upon cholesterol depletion from the plasma membrane ^23,24^.

The model predicts the lipid monolayer fragments facing the caveolin discs to be enriched in cholesterol, whereas the membrane pieces of the bilayer fragments would have lowered cholesterol concentrations (Fig.4). Since the lengths of the optimal gaps between the discs constitute several nanometers ^10,13,29,30^ and, thus, are considerably smaller than the 14nm large discs, the distal monolayer of the system is predicted to have an overall elevated cholesterol content which is in accord with observations ^66^.

It has to be emphasized that, within the proposed mechanism, the effects of cholesterol are of mechanical rather than chemical origin and are related to the negative intrinsic curvature, *ζ*, of cholesterol molecules ^32^. Therefore, similar effects on the membrane shaping by caveolin discs can be produced by other lipids with strongly negative intrinsic curvatures such as, e.g., diacylglycerols ^67^. Interestingly, specific diacylglycerol species were shown to be enriched in h-caveolae generated in *E*.*coli* (Walser 2012) and our model would predict a role for these or related lipids in membrane sculpting by caveolin in that system.

### Model assumptions

One limitation of the presented model is its two-rather than three-dimensional character. This consideration substantially simplified the analysis allowing us to obtain analytical and easily understandable relationships between the geometrical features of the system and the experimentally relevant parameters such as the cholesterol mole fraction, *c*_0_. Yet a shortcoming of this simplification is that a comparison between the predicted membrane conformations and the ones observed in experiments can have only a semi-quantitative meaning.

In our computations, we assumed a vanishing geometrical curvature of the bilayer fragments flanking the disc part and consisting of the monolayers II and III. This can now be justified *a posteriori* based on the obtained results. The bilayer curvature would be generated if there were an asymmetry between the monolayers II and III. This asymmetry can be a consequence of a difference between the monolayers in either the deformations of tilt-splay, or the cholesterol concentration. According to the obtained results, in the energetically optimal state of the system, both the tilt-splay and the cholesterol concentration are equal in the monolayers II and III and, hence, there is no asymmetry between the leaflets.

We used simplifying assumptions concerning the membrane lipid composition. We considered the membrane as consisting of two lipid species, one lipid with a vanishing and the second, mimicking cholesterol, with a strongly negative molecular intrinsic curvature, *ζ*. In reality, a cell plasma membrane contains a variety of lipid species with different non-vanishing intrinsic curvatures, all of which would exhibit some extent of redistribution within the system according to the same mechanism as considered above for cholesterol. Consideration of the redistribution of additional lipids would predict even larger effective curvatures generated by caveolin discs.

Finally, the model does not specify the interaction between the lipid molecules and the caveolin discs. While particular chemical interactions between lipids and the caveolar coat proteins have been proposed ^18,25,26^ our model describes them by a general parameter of the difference in contact energies, Δ*ε*, whose value was assumed to have an order of 1*k*_*B*_*T*. Verification of this assumption requires detailed quantitative information on the energies of lipid-caveolin interactions that is currently unavailable.

## Acknowledgements

MMK acknowledges support from the Israel Science Foundation (Grant No. 1994/22) and holds the Joseph Klafter Chair in Biophysics. RGP was supported by an Australian Research Council (ARC) Laureate Fellowship (FL210100107).

## Appendix A

### Equations of equilibrium

The partial minimization of the system’s energy can be performed by keeping constant values of the boundary tilt, *ϕ*_*B*_, and kink, *φ*, and seeking, for each system’s monolayer, for the distributions of the tilt angle, *ϕ*(*x*), splay,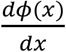, and cholesterol molar ratio, *c*(*x*), which minimize the monolayer’s energy. This boils down to minimization for each monolayer of the sum of the monolayer’s elastic, *F*_*el*_ and cholesterol distribution, *F*_*chol*_, energies,

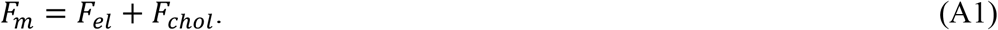

According to (Eqs.6,7,10) and definition of the multiplier *D* given in the main text, the area density of this energy can be presented as

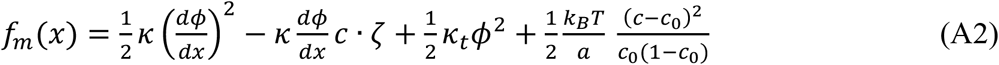

so that the monolayer energy is

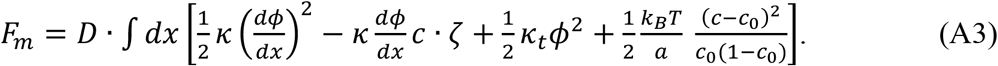

The conditions of the energy minimum are expressed by two Lagrange equations ^68^, which corresponds to the vanishing variation of the energy, δ*F*_*m*_ = 0, upon variations of the functions, δ*c*(*x*), and, δ*ϕ*(*x*), and have a meaning of the equations of equilibrium.

The equation of resulting from the variation of the cholesterol mole fraction, δ*c*(*x*), relates the cholesterol distribution and the splay

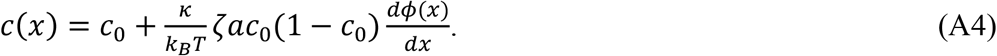

Using (Eq.A4), the monolayer energy per unit area (Eq.A2) takes the form,

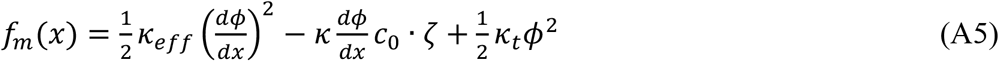

where

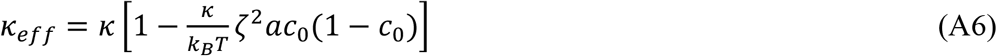

is the effective bending (splay) modulus of the monolayer.

The total monolayer energy can be obtained by integration of the energy density (Eq.A5) over the monolayer are

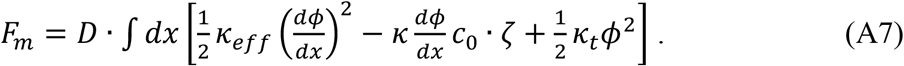

The second equation of equilibrium that corresponds to variation of the tilt angle, δ*ϕ*(*x*), and fixed splay at the monolayer boundaries ((Eqs.2,3) of the main text) has the form,

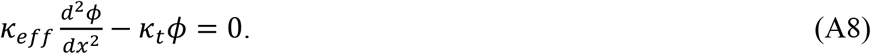

### Tilt, splay and cholesterol distributions in the system’s monolayers

Solution of the equilibrium equation (Eq.A8) satisfying the boundary conditions (Eqs.2,3 of the main text) for each monolayer of the system gives the distribution of the tilt angle, *ϕ*(*x*), and, hence, the splay,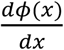. Substitution of the obtained splay function into (Eq.A4) gives the expression for the cholesterol mole fraction, *c*(*x*).

The coordinate axis *x* for each s chosen directed from the external to the internal monolayer boundary with the origin (*x* = 0) at the former.

Based on (Eq.A8), we define the characteristic decay length of the tilt deformation,

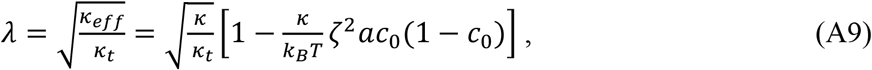

referred to in the main text as the operational length scale.

Using (Eq.A9) we define the dimensionless coordinate, 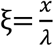, the dimensionless radius of the caveolin disc, 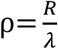, and the dimensionless half distance between the discs, 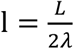.

For the monolayer I, we obtain for the tilt and splay,

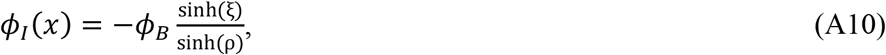

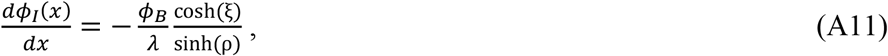

for the mole fraction of cholesterol,

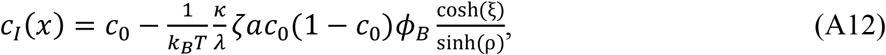

and for the area average of the mole fraction of cholesterol,

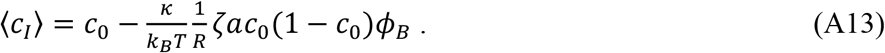

For the monolayer II we obtain,

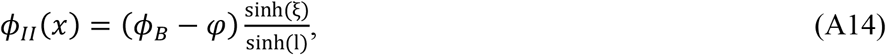

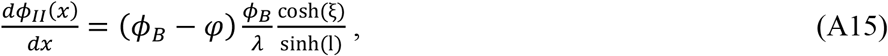

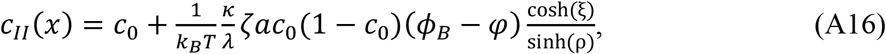

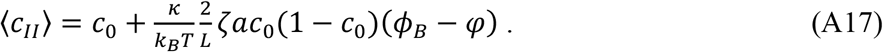

For the monolayer III we obtain,

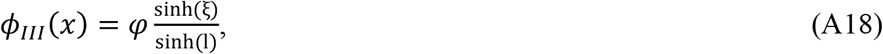

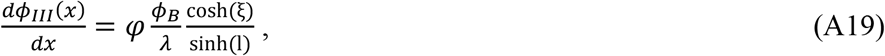

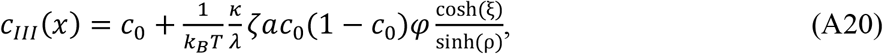

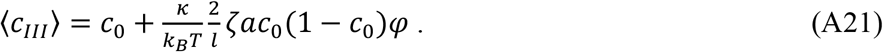

### The monolayer energies

Using the obtained expressions for the distributions of tilt, splay, and cholesterol mole fraction along the monolayer plains, we determine for each monolayer the area density of the monolayer energy, *f*_*m*_(*x*) (Eq.A5). Integrating the obtained *f*_*m*_(*x*) over the monolayer area, which corresponds to integration from *x* = 0 to *x* = *R* for the monolayer I, and from *x* = 0 to *x* = *L*/2 for the monolayers II and III, we get the monolayer energies, *F*_*m*_(Eq. A7).

For the monolayer I we obtain,

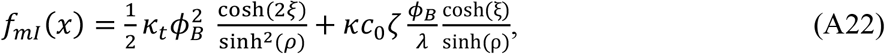

and

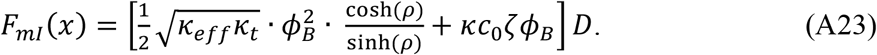

For the monolayer II we obtain,

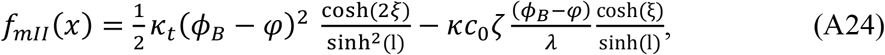

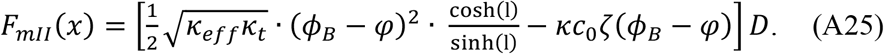

For the monolayer III we obtain,

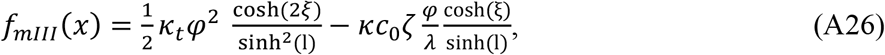

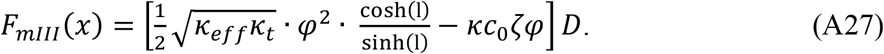

### The total energy of the system

Using the above definition (Eq.A1) of the monolayer energy, *F*_*m*_, the total energy of the system, *F* ((Eq.4) of the main part) can be presented

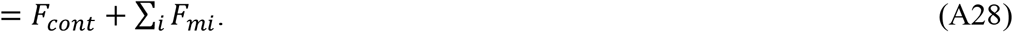

Using (Eq.5) of the main text for the contact energy, *F*_*ccct*_, and (Eqs.A23,A25,A27) for the monolayer energies, we obtain for the total energy

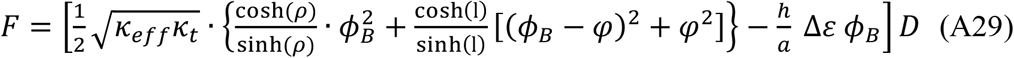

### Minimal energy state of the system and interaction between the caveolin discs

The minimal energy state of the system corresponding to the given distance, *L*, between the discs, is determined by minimization of the energy, *F* (Eq.29), with respect to the boundary tilt, *ϕ*_*B*_, and kink, *φ*, angles. This results in the optimal values of these angles,

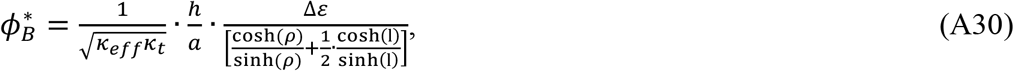

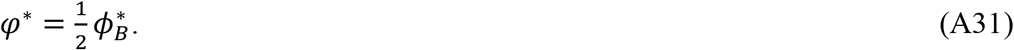

While the system’s configuration has a faceted appearance (Fig.1D,E of the main part), it can be described by an effective curvature, *J*_*eff*_, that is expressed through the kink angle, *φ*^∗^, the half distance between the discs, *R* + *L*/2, and the disc radius, *R*, by

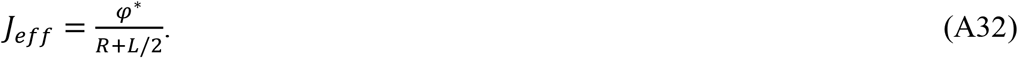

The minimal energy of the system is

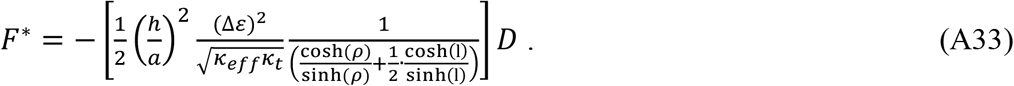

The force of interaction, ℱ between adjacent caveolin discs is given by

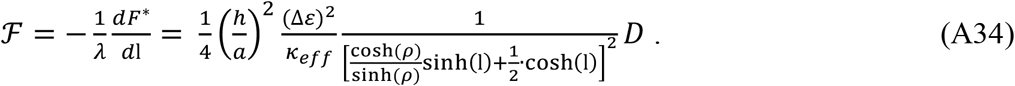

